# Advancing Inter-brain Synchrony Measurement: A Comparative Hyperscanning Study of High-Density Diffuse Optical Tomography and Functional Near-infrared Spectroscopy

**DOI:** 10.1101/2025.07.02.662877

**Authors:** Shuo Guan, Yuhang Li, Yingbo Geng, Dongyun Li, Qiong Xu, Dalin Yang, Yingchun Zhang, Rihui Li

## Abstract

Inter-brain synchrony (IBS), measured by hyperscanning, refers to the synchronization of multiple individuals’ brain activities during social interactions. Traditional fNIRS-based hyperscanning suffers shortcomings like low spatial resolution and high susceptibility to superficial interference, causing imprecise estimation of IBS in complex social tasks. This study aims to fill the knowledge gap by comprehensively assessing how high-density diffuse optical tomography (HD-DOT), an enhanced alternative to fNIRS, can benefit hyperscanning studies of complex social interactions. Sixteen dyads were engaged in both collaborative and individual tangram puzzle tasks, and their brain activities were recorded simultaneously using HD-DOT and fNIRS. We found that HD-DOT demonstrated significantly stronger IBS and identified more brain regions with significant IBS compared to fNIRS during the collaborative task. Specifically, while fNIRS detected IBS only in the dorsolateral prefrontal cortex (DLPFC) and supramarginal gyrus (SMG), HD-DOT revealed additional IBS in the superior temporal gyrus (STG). Additionally, compared to the individual task, the collaborative task showed increased IBS in HD-DOT, not only in the DLPFC but also in the SMG, frontal eye fields (FEF), and inferior frontal gyrus (IFG). By highlighting the superior spatial resolution and sensitivity of HD-DOT in capturing detailed and extensive neural activity during complex social interactions, our findings for the first time clarified the potential strengths of HD-DOT in measuring IBS over traditional fNIRS. These advances provide a stronger empirical foundation for investigating the neural basis of social interaction, paving the way for future research on real-world, dynamic group behaviors.

**Highlights:**

- First application of HD-DOT for inter-brain synchrony measurement.
- First demonstration of HD-DOT’s superiority over fNIRS in IBS measurement.
- HD-DOT detects stronger IBS and more brain regions than fNIRS.

## 1. Introduction

Human social interaction fosters complex problem-solving and creativity, serving as a key driver of economic growth and social progress (Balconi et al., 2017; Tei et al., 2017). Understanding how individuals engage with one another during social exchanges, namely interpersonal interaction, is not only essential for unravelling our social nature but also provides critical insights into the neural underpinnings of social-deficit disorders such as autism spectrum disorder (R. Li et al., 2021).

A promising framework for studying interpersonal interaction is inter-brain synchronization (IBS), the phenomenon in which brain activities synchronize across individuals during social exchanges (Hu et al., 2017; Liu et al., 2023). Accumulating evidence suggests that IBS plays a fundamental role in the cognitive and emotional processes underlying complex social behaviors, particularly in collaborative tasks where individuals work toward shared goals(Nguyen et al., 2020; Yoshioka et al., 2021).

The emergence of hyperscanning technology has enabled the simultaneous recording of brain activity for calculating IBS among multiple interacting individuals. Among neuroimaging tools, functional near-infrared spectroscopy (fNIRS) has emerged as a leading hyperscanning approach for measuring IBS during naturalistic social interactions due to its high mobility and tolerance to movement (Cui et al., 2012; Quaresima & Ferrari, 2019). To date, fNIRS-based hyperscanning studies have examined IBS across diverse social contexts, including parent-child communication, teacher-student learning, and team cooperation/competition (Nguyen et al., 2021; Sun et al., 2021; Wu et al., 2025). These studies reported significant IBS modulations in key brain regions of interacting dyads, such as the dorsolateral prefrontal cortex (DLPFC) and the temporoparietal junction (TPJ) (Donaldson et al., 2015; Hakim et al., 2023; Nejati et al., 2023). Yet, traditional fNIRS faces notable limitations, particularly in spatial resolution and susceptibility to superficial contamination (Scholkmann et al., 2014). These constraints can impede the precise localization of brain activity during complex social tasks, potentially obscuring subtle but important neural patterns that manifest during social interactions.

To address the inherent methodological challenges, high-density diffuse optical tomography (HD-DOT) is a promising alternative to fNIRS, offering enhanced spatial resolution and improved depth sensitivity (Eggebrecht et al., 2014; Fan et al., 2025; Liao, 2012).HD-DOT represents a significant advancement in optical neuroimaging, which comes with dense arrays of optical sensors and sophisticated image reconstruction algorithms (Ayaz et al., 2022; Vidal-Rosas et al., 2023). This technology enables the creation of three-dimensional reconstructions of brain hemodynamics, providing more detailed and accurate representations of neural activity (Tripathy et al., 2024; Zeff et al., 2007). Despite the growing evidence supporting the superior spatial resolution over fNIRS, whether and how the improved spatial resolution and reduced susceptibility to superficial contamination of HD-DOT would benefit hyperscanning studies in investigating complex social interaction remains unknown.

To address this gap, the present study conducted a comparison study using both traditional fNIRS and HD-DOT techniques during a hyperscanning-based social interaction task. Specifically, we designed experiments where participants engaged in collaborative and individual tangram puzzle tasks, while their brain activities were simultaneously recorded using both HD-DOT and fNIRS montage designs. By comparing the performance from these two imaging modalities, we aimed to validate the strengths of HD-DOT over fNIRS in measuring IBS, thereby providing evolved approaches for understanding the neural basis of human social interaction.

## 2. Methods

### 2.1 Participants

38 healthy volunteers (16 females, 22 males, aged 24.4 ± 4.7 years) were randomly assigned to participate in this study. None of the participants reported a history of significant medical, neurologic, or psychiatric illness. All participants were right-handed, already familiar with the game of tangram puzzle, and spoke Mandarin as their primary language. All study protocols were conducted following the Declaration of Helsinki and were approved by the Institutional Review Boards at the University of Macau. Informed consent was obtained from each participant before the study.

### 2.2 Experimental paradigm

Before the initiation of the experiment, the participants received a comprehensive briefing on the rules and procedures. The experimental paradigm consisted of three parts and lasted approximately 20 minutes (Figure 1A). First, each dyad engaged in a 6-minute cooperative tangram puzzle task, during which they were required to communicate continuously and complete as many puzzles as possible (Figure 1B). This was followed by a 5-minute resting period. Afterward, each participant completed the same tangram puzzle individually, without any verbal communication or eye contact with their partner (Figure 1C). Prior to the experiment, participants were given sufficient practice time to become acquainted with the tangram puzzles. The number of puzzle pieces completed by each participant and dyad was recorded. Following the experiment, participants were asked to rate the difficulty of both tasks on a scale of 1 to 10.

**Figure 1.**
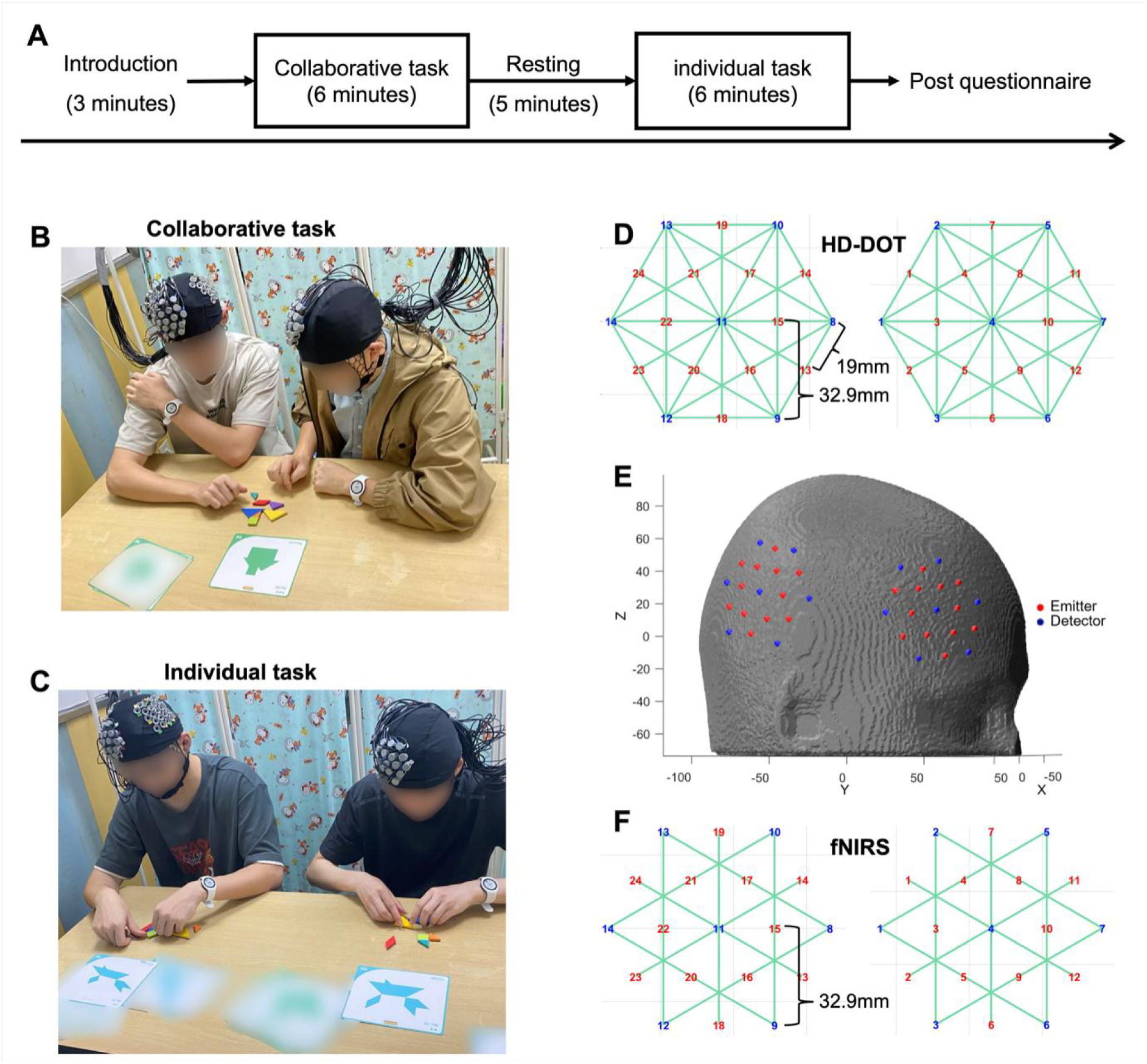
Experimental design. (**A**) The paradigm included collaborative and individual tangram puzzle tasks, along with an introduction session, rest period, and questionnaire process. (**B**) Participants communicated verbally and made eye contact while jointly working on one puzzle in the collaborative task. **(C**) Participants worked independently on identical puzzles without communication in the individual task. **(E**) Sensor distribution, including both the HD-DOT and traditional fNIRS, was designed to cover the right prefrontal region and part of the temporal and parietal area. Channel arrangement of HD-DOT **(D**) and fNIRS **(F**).

### 2.3 fNIRS and HD-DOT acquisition

A fNIRS system (Danyang Huichuang, China) was used to conduct traditional fNIRS and HD-DOT recordings. The device utilized continuous-wave measurements at two wavelengths (750 and 835 nm) to measure the hemodynamic activity of the targeted right prefrontal cortex and temporoparietal at a sampling frequency of 11 Hz. Specifically, 24 emitters and 14 detectors (Figure 1E) were used to form 84 multi-distance channels for HD-DOT recording on the scalp (Figure 1D) as well as 36 regular channels for traditional fNIRS recording (Figure 1F). The design of the HD-DOT montage was inspired by the research of Gao et al., and it adopted a hexagonal pattern layout (Gao et al., 2023). The short-distance detectors were placed at 19 mm from the light sources. These were mainly used to detect the optical signals from superficial tissues such as the scalp and skull (Von Lühmann et al., 2024). The long-distance detectors were set at 32.9 mm from the light sources, with the primary function of obtaining the functional signals from deep-brain tissues, reflecting the changes in hemoglobin concentration caused by neuronal activities. All the optodes of fNIRS and HD-DOT recordings was mounted by a 3-D printing cap designed by the Atlasviewer software (Aasted et al., 2015).

### 2.4 Data quality assessment and preprocessing

We applied standardized preprocessing procedures to both fNIRS and HD-DOT data, as shown in Figure 2. In terms of quality assessment, three metrics were obtained through the processing pipeline in NeuroDOT (https:// www.nitrc.org/projects/neuroHD-DOT). Firstly, the pulse signal-to-noise ratio (SNR) was calculated by determining the ratio of signal power within the 0.5–2 Hz band to the bandwidth-scaled median noise power in adjacent frequency ranges across the measurement cap. This metric assessed the system’s sensitivity to vascular physiology. A threshold SNR of 1 dB was set to exclude experimental runs with poor optode-scalp coupling (Yang et al., 2024). Secondly, the good measurement (GM) percentage was defined as the proportion of measurements with a temporal standard deviation less than 7.5%. To ensure consistent spatial coverage of the field of view, runs with less than 80% GM were excluded from further analysis (Yang et al., 2025). Thirdly, the global variance of the temporal derivative (GVTD) was used to quantify the levels of motion. An empirical GVTD threshold of 0.1% was applied to remove data with high levels of motion (Di Sciacca et al., 2022; Sherafati et al., 2020).

**Figure 2.**
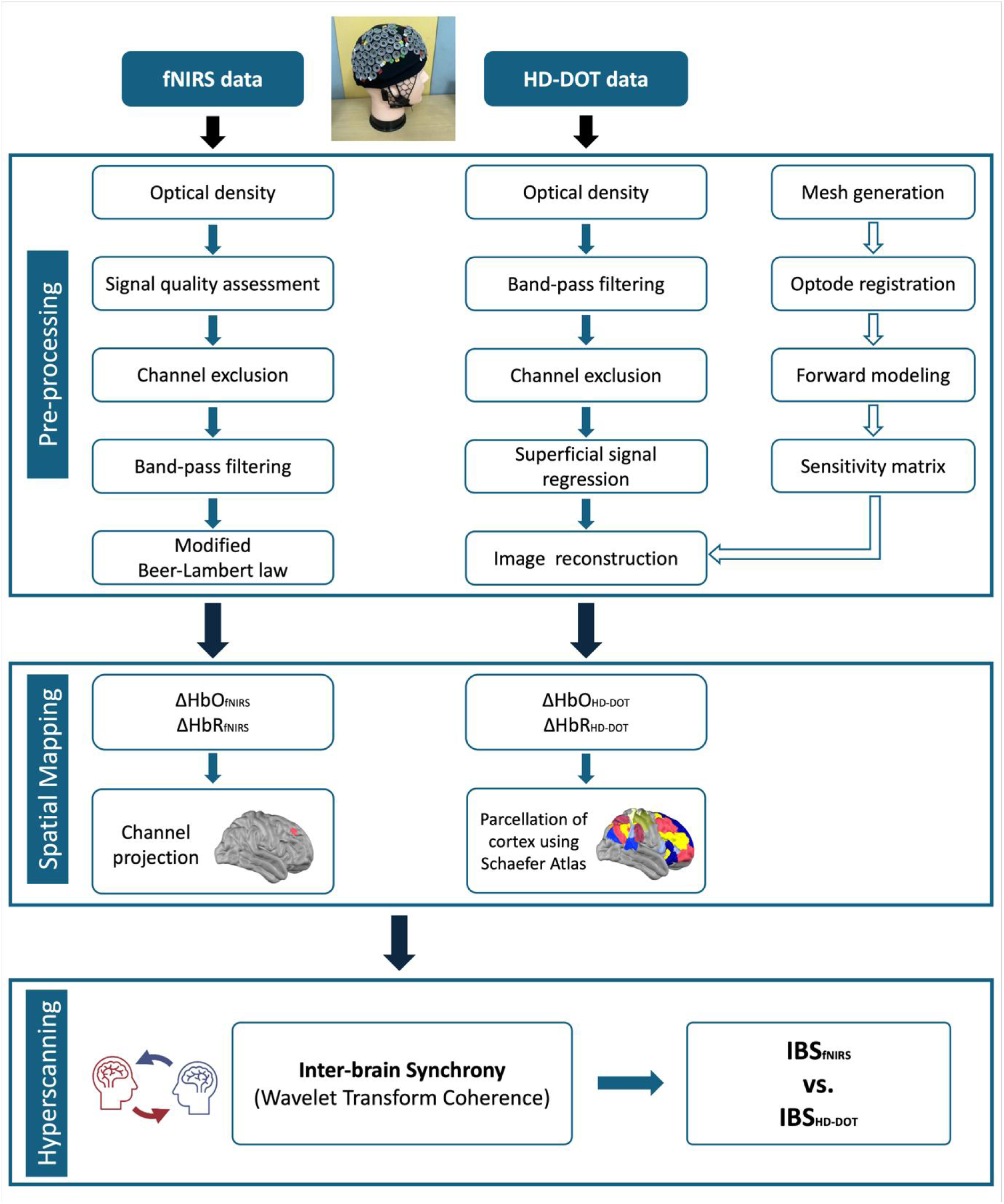
The processing pipelines of fNIRS and HD-DOT, including preprocessing, image reconstruction of HD-DOT, spatial mapping, and hyperscanning analysis.

After data quality assessment, high-pass filtering was performed with a 0.02 Hz cutoff to reduce long-term drift, and low-pass filtering with a cutoff frequency of 2 Hz was also carried out to remove high-frequency noise (Hassanpour et al., 2014). Compared with fNIRS, HD-DOT had an additional step called superficial signal regression, which was achieved by averaging all short-distance measurements to estimate global systemic signals and linearly regressing these out of every source-detector time trace.

### 2.5 Forward modeling and Image reconstruction

For the forward modeling, we used a heterogeneous head model composed of five tissue types (i.e., scalp, skull, grey matter, white matter, and cerebrospinal fluid), each with specific optical properties (Ferradal et al., 2014). A finite-element, wavelength-dependent forward light-model was employed to calculate the sensitivity for the full array of source-detector measurements. The non-linear ICBM152 atlas from the Montreal Neurological Institute (MNI) was used as the anatomical model, and the sensitivity matrix was created based on the atlas and optode positions using NIRFASTer (Dehghani, Eames, et al., 2009). The sensitivity matrix was inverted using a spatially variant regularization parameter of 0.1, a Tikhonov regularization parameter of 0.01, and a 3-mm full-width at half-maximum Gaussian 3D smoothing kernel to remove speckly noise (Dehghani, Srinivasan, et al., 2009). The inverted matrix was used to reconstruct absorption coefficient changes for each wavelength. Relative changes in the concentrations of oxygenated (HbO), deoxygenated (HbR), and total hemoglobin (HbT) were obtained from the absorption coefficient changes by the spectral decomposition of the extinction coefficients of HbO and HbR at these two wavelengths (Hassanpour et al., 2014).

### 2.6 Cortical parcellation

The reconstructed signals were parcellated into distinct anatomical regions to enable direct cross-participant comparisons following the Schaefer atlas (Schaefer et al., 2018), yielding a total of 118 valid regions of interest (ROIs). The Nilearn toolbox was used to compare the performance of fNIRS and HD-DOT data within the same spatial coordinate system (https://www.nitrc.org/projects/nilearn/). Specifically, we used this toolbox to identify the corresponding ROIs in the parcellation template (the same template used for HD-DOT segmentation) for the MNI coordinates of the fNIRS channels. This approach streamlined the process of comparing fNIRS and HD-DOT data, ensuring that the analysis is conducted within a unified and consistent framework.

### 2.7 Inter-brain synchrony

The Wavelet Transform Coherence (WTC) method was used to assess the IBS between each dyad for both the fNIRS and HD-DOT data (Cui et al., 2012). Specifically, for each pair of time series (channel signal for fNIRS and ROI signal for HD-DOT), we computed the WTC matrices using the wavelet toolbox in MATLAB 2023a (Grinsted et al., 2004). The WTC matrices (frequency × time) were then averaged across time points and Fisher-to-z transformed for each task condition as well as the resting state. We then applied a data-driven method to select the frequency of interest by comparing the differences in the WTC values across all pairwise combinations of conditions (Hu et al., 2021). The selected frequency band excluded high or low frequency noise, such as physiological noise related to blood pressure (approximately 0.1 Hz), respiration (approximately 0.2–0.3 Hz), and heart rate (1 Hz). Finally, the WTC values at selected frequency bands were averaged for each channel and ROI pair.

### 2.8 Statistical analysis

A series of paired *t*-tests were applied to identify the task-evoked IBS for both fNIRS and HD-DOT data. Specifically, comparisons were conducted between the collaborative task and the resting state, the collaborative task and the individual task, and the individual task and the resting state, respectively. The resulting *p*-values of all ROI comparisons were corrected using the false discovery rate (FDR) with the Benjamini-Hochberg approach (*p* < 0.05). We also used a series of paired *t*-tests to assess the strength of the IBS values at the identified ROI between the HD-DOT and fNIRS modalities. Correlational analyses were conducted to assess the relationship between the IBS at the identified ROI and behavioral performance. The same FDR method was also applied to correct the multiple comparison effect.

## 3. Results

### 3.1 Demographic characteristics and task performance

As shown in Table 1, 32 subjects (13 females, 19 males) completed the entire experiment. Three dyads were excluded from all analyses due to incorrect task completion or substandard signal quality. The mean self-assessed difficulty of collaborative task was 5.62 ± 1.92, while that of the individual task was 6.03 ± 2.07. There was no significant difference between the two task types (t = −0.34, *p* = 0.36). The mean number of puzzles solved in collaborative tasks was 3.31 ± 1.86, and for individual tasks, it was 2.34 ± 1.67. There was a significant difference between the two task types (t = 2.44, *p* = 0.02).

**Table 1.** Behavior performance. The number of puzzles solved by the participants and the self-assessed task difficulty in the collaborative and individual puzzle tasks.

| Subject | Age | Collaborative task |  | Individual task |  |
| --- | --- | --- | --- | --- | --- |
|  |  | Number of completed puzzles | Self-assessed difficulty | Number of completed puzzles | Self-assessed difficulty |
| 16 dyads | $24.03 \pm 4.56$ | $3.31 \pm 1.86$ | $5.63 \pm 1.92$ | $2.34 \pm 1.67$ | $6.03 \pm 2.07$ |

### 3.2 Inter-brain synchrony during collaborative task

We first examined whether significant IBS occurred during the collaborative task as compared to the resting state. HD-DOT-based analysis revealed enhanced IBS in six ROI pairs during the collaborative task, including the dorsolateral prefrontal cortex (DLPFC), as well as the supramarginal gyrus (SMG) and superior temporal gyrus (STG), which were situated within the temporoparietal junction (TPJ) (Figure 3A). For the fNIRS, significantly enhanced IBS was observed in the DLPFC and SMG regions (Figure 3B). There was an overlap in the regions identified by both HD-DOT and fNIRS, including the DLPFC and SMG. In particular, the computed IBS strength at the two overlapping regions was significantly stronger in the HD-DOT compared to the fNIRS data (Figure 3C, DLPFC, t = 4.20, *p* < 0.01; Figure 3D, SMG, t = 2.29, *p* = 0.04, FDR corrected).

**Figure 3.**
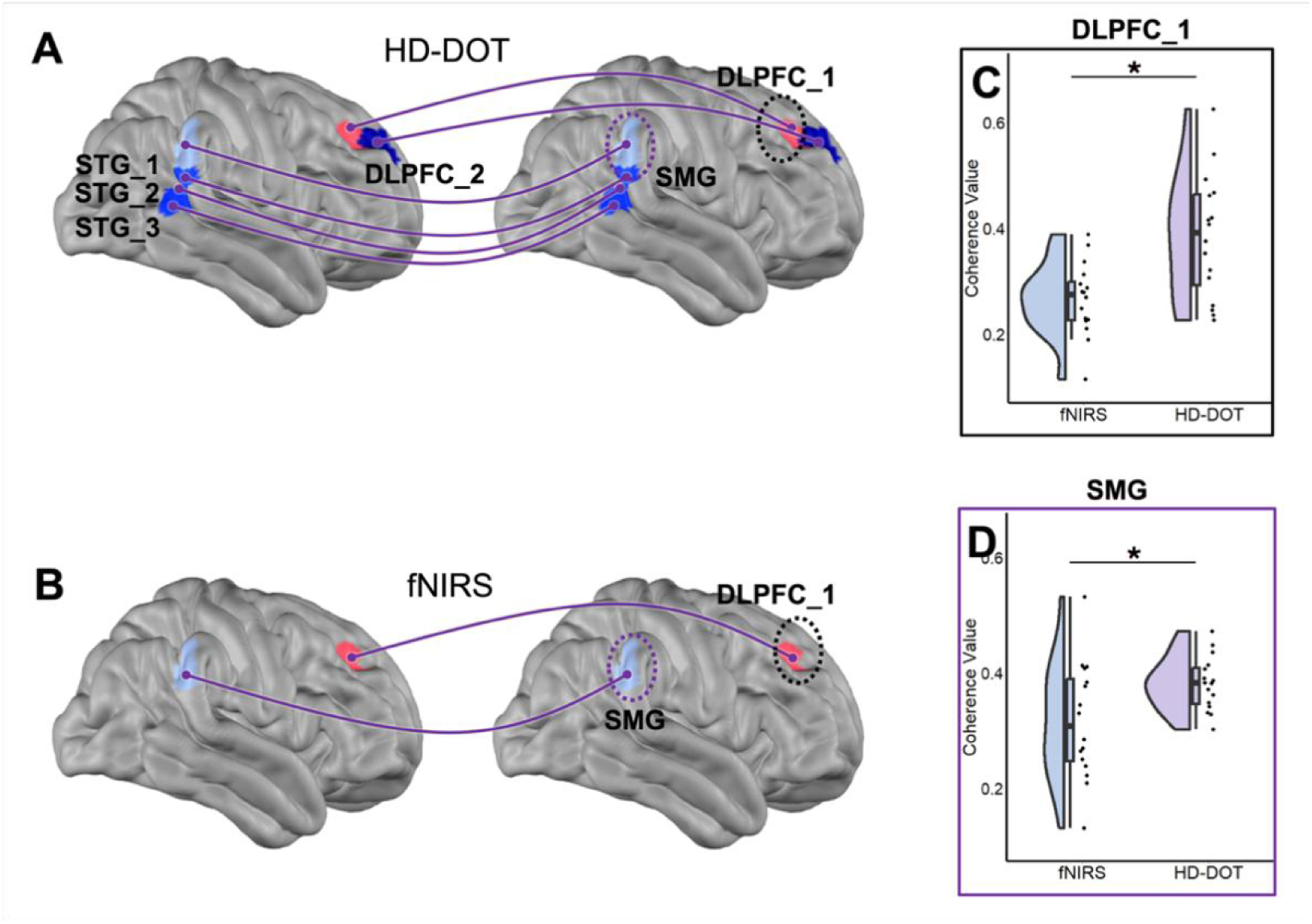
The IBS pattern during the collaborative task versus the resting state. (**A**) Significant IBS in six ROIs were derived from the HD-DOT data, including DLPFC_1, DPLFC_2, SMG, STG_1, STG_2, and STG_3. **(B**) Significant IBS in two ROIs were derived from fNIRS including DLPFC_1 and SMG. **(C-D**) The comparisons of IBS in DLPFC and SMG between the HD-DOT and fNIRS approaches. (\**p* < 0.05, FDR corrected).

We further investigated whether significant IBS occurred during the collaborative task compared to the individual task. During the collaborative task, there was a significantly enhanced IBS identified by HD-DOT in four specific ROI pairs, including the DLPFC, SMG, frontal eye fields (FEF), and inferior frontal gyrus (IFG) (Figure 4A). For the fNIRS data, significantly enhanced IBS was observed only in the DLPFC region (Figure 4B). Similarly, significantly stronger IBS strength in the overlap ROI (i.e., DLPFC) was observed from the HD-DOT data compared to the fNIRS data (Figure 4C, t = 2.53, *p* = 0.04, FDR corrected).

**Figure 4.**
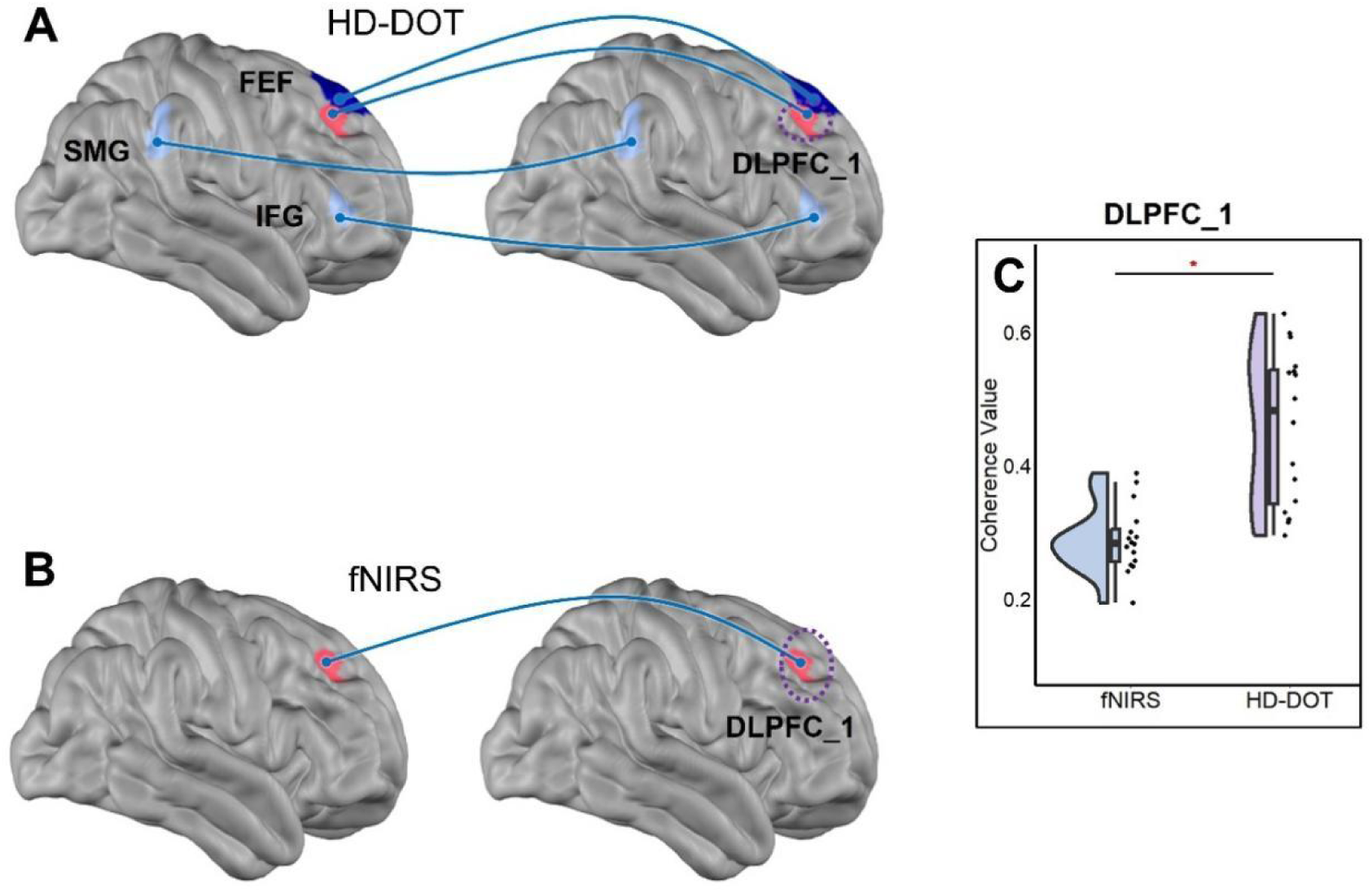
The IBS pattern during the collaborative task versus the individual task. (A) Significant IBS in four ROIs were derived from HD-DOT data, including DLPFC_1, FEF, SMG, and IFG. (B) Significant IBS in DLPFC_1 was derived from fNIRS. (C) The comparisons of IBS in DLPFC_1 between the HD-DOT and fNIRS approaches. (\**p* < 0.05, FDR corrected).

### 3.3 Inter-brain synchrony during individual task

As an exploratory analysis, we also examined the significant IBS during the individual puzzle task relative to the resting state. Significantly enhanced IBS in the angular gyrus (AG), located within the TPJ, was identified across dyads from the HD-DOT data (Figure 5A). Comparatively, the fNIRS data also revealed significantly enhanced IBS in the AG (Figure 5B). However, HD-DOT identified a larger activation area and stronger IBS strength in the AG compared to the fNIRS data (t = 3.21, *p* = 0.02, FDR corrected) (Figure 5C).

**Figure 5.**
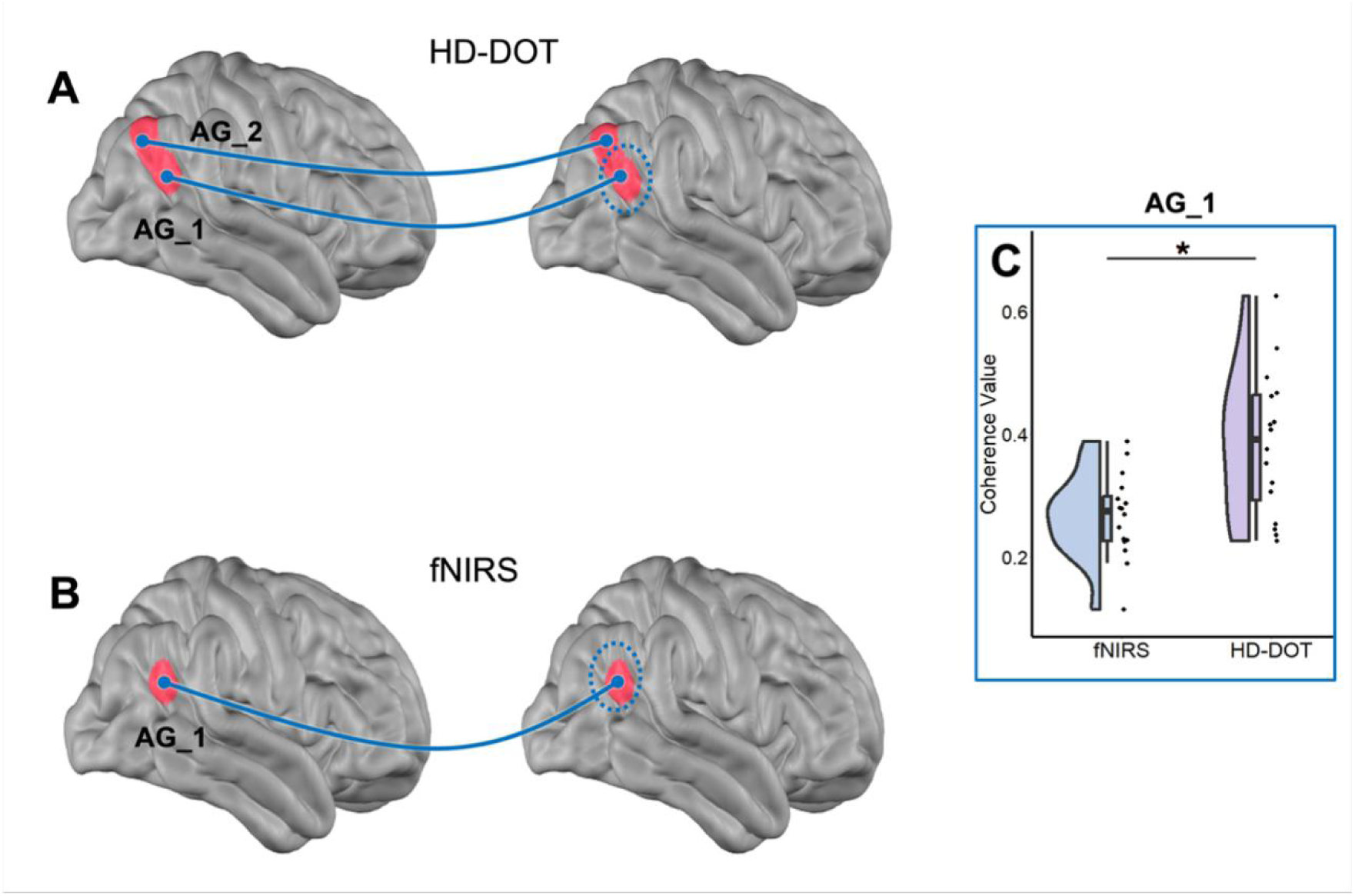
The IBS pattern during the individual task versus the resting state. (A) Significant IBS in two ROIs were derived from HD-DOT data including AG_1 and AG_2. (B) Significant IBS in AG_1 was derived from fNIRS. (C) The comparisons of IBS in AG_1 between the HD-DOT and fNIRS approaches. (\**p* < 0.05, FDR corrected).

### 3.4 Task performance related to the correlation with IBS

We employed Pearson correlation analysis to investigate the relationship between the task performance (i.e., number of completed tangram puzzles) and the IBS in identified brain regions for both HD-DOT and fNIRS. As shown in Figure 6, in the HD-DOT data, there was a significant positive correlation between IBS in DLPFC and the number of completed puzzles (r = 0.506, *p* = 0.045). However, in the fNIRS data, while a positive correlation trend was observed between IBS and the number of completed puzzles, this correlation was not statistically significant (r = 0.192, *p* = 0.476).

**Figure 6.**
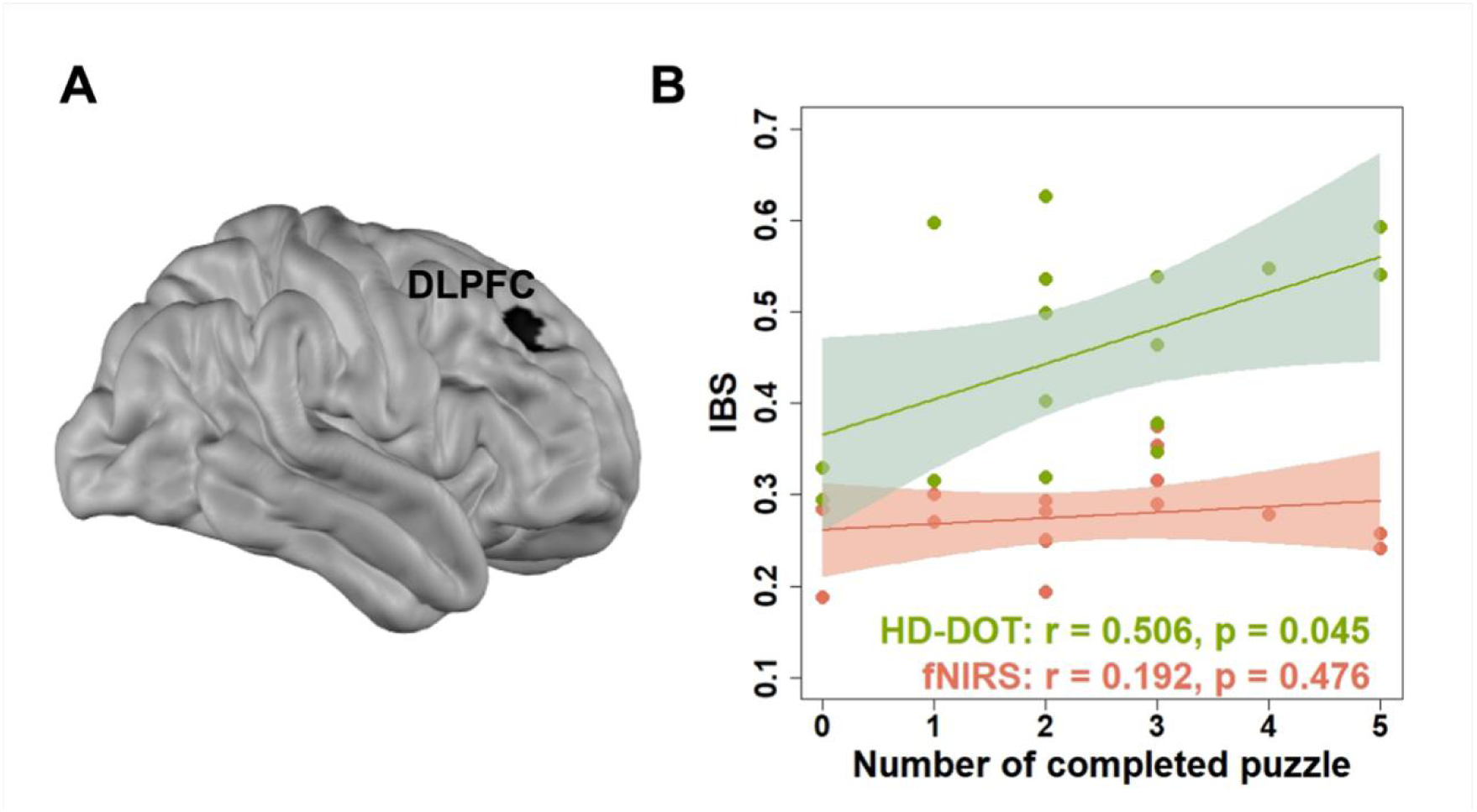
(**A**) The dyadic IBS in DLPFC was identified by both the HD-DOT and fNIRS. **(B**) HD-DOT hyperscanning analysis revealed a significant positive correlation between the IBS and the dyadic behavioral performance.

## 4. Discussion

fNIRS has become a routine technique for hyperscanning studies in social science and neuroscience due to its high ecological validity. Yet, conventional fNIRS is subject to low spatial resolution and highly susceptible to superficial contamination, limiting the reliability and reproducibility of scientific findings. The present study directly compared HD-DOT, the enhanced form of fNIRS, and traditional fNIRS in a hyperscanning paradigm to evaluate their efficacy in capturing IBS during collaborative interactions. Our results demonstrated that HD-DOT outperformed traditional fNIRS in detecting IBS across key brain regions and linking dyadic IBS to behavioral performance, underscoring its potential as a transformative tool for social neuroscience.

During the collaborative task, both fNIRS and HD-DOT identified significant IBS within the DLPFC and TPJ region (e.g., SMG and STG) compared to the resting state. The IBS at such overlapping regions aligns with a growing body of research underscoring the DLPFC’s critical role in executive functions, working memory, and cognitive control during social interaction (Frith & Singer, 2008; Miller & Cohen, 2001). The enhanced IBS in the DLPFC may reflect the coordination of higher-order cognitive processes necessary for shared intentionality and goal-directed collaboration (Diwadkar et al., 2000). The TPJ region such as SMG and STG, implicated in effective information sharing and joint decision-making, may facilitate the integration of individual perspectives toward a common solution (Hilgenstock et al., 2014).

The present study stands as the first study to validate the hypothesis that HD-DOT significantly enhances the detection and localization of IBS compared to traditional fNIRS in collaborative tasks. While both modalities identified IBS in core social cognition regions (e.g., DLPFC, SMG), HD-DOT uniquely captured IBS in broader cortical regions (STG, IFG) linked to social communication and further demonstrated stronger IBS-behavior correlations. The STG absent in fNIRS but identified by HD-DOT, is associated with theory of mind and perspective-taking (Saxe & Kanwisher, 2003), suggesting its involvement in understanding the intentions and beliefs of the co-actor during collaborative puzzle-solving. This observation is consistent with previous fMRI studies reporting similar activations during tasks requiring communication and joint attention, further supporting the validity of HD-DOT findings (Wang et al., 2022; H. Xie et al., 2020). Our findings also align with recent HD-DOT validation studies in individual neuroimaging where the DOT model with short-separation channels significantly improved the fNIRS image reconstruction quality (Gao et al., 2023), extending their implications to dyadic interaction paradigms. Taking these together, the methodological advancement of HD-DOT addresses two critical limitations of conventional fNIRS. First, HD-DOT improved depth sensitivity through multi-distance measurements, which effectively distinguishes superficial hemodynamic artifacts from true cortical activity (Chiarelli et al., 2016). Secondly, HD-DOT enhanced spatial resolution by dense optode arrays and 3D tomographic reconstruction, allowing for more precise localization of brain activity.

The expanded IBS patterns detected by HD-DOT offer new insights into the distributed neural architecture supporting collaborative cognition. Particularly, HD-DOT revealed additional IBS activity in the broader TPJ, FEF, and IFG during the collaborative task (versus individual task). These regions are implicated in joint attention, language processing, and action coordination (D. Xie & Xue, 2025), suggesting that HD-DOT captures a broader neural signature of social interaction. Particularly, HD-DOT uniquely captured significant IBS in IFG, an IBS finding absent in fNIRS data, demonstrating its unique capacity to monitor the complex cognitive processes related to social interaction. In the context of collaborative tasks, the IFG enables individuals to interpret their partners’ intentions, synchronize actions, and process both semantic and pragmatic aspects of communicative language (Fishburn et al., 2018; Y. Li et al., 2021). The absence of these findings in fNIRS data may explain historical inconsistencies in social neuroscience literature (Czeszumski et al., 2022; Nam et al., 2020), where superficial signal contamination could obscure temporally coordinated activity in broader and deeper brain regions.

Beyond identifying IBS at additional brain regions, HD-DOT demonstrated superior detection power even in brain regions where both modalities showed significant IBS. The significantly stronger IBS strengths in DLPFC and SMG likely stem from HD-DOT’s enhanced SNR through the superficial signal regression, which effectively removes scalp hemodynamic contamination that typically inflates between-subject variance in fNIRS (Guan et al., 2025). In addition, the high-density sampling through multi-distance probes enables more accurate partial volume correction, particularly critical in prefrontal regions with complex cortical folding patterns (Eggebrecht et al., 2014). Our empirical findings here demonstrate HD-DOT’s superior depth discrimination, particularly in anterior brain regions, suggesting historical fNIRS hyperscanning studies might have systematically underestimated prefrontal contributions to social coordination.

Another key finding of our study is the discrepancy in the strength of the relationship between IBS in DLPFC and task performance as measured by HD-DOT and fNIRS. The stronger IBS-performance correlation observed in HD-DOT highlights its superior ecological validity for hyperscanning research. This finding aligns with theoretical frameworks positing that dyadic prefrontal synchronization facilitates cooperative success by enhancing mutual predictability and reducing cognitive load (Broadbent et al., 2023). Conversely, the lack of a significant correlation in fNIRS-derived IBS suggests that superficial contamination or spatial inaccuracy may have hindered the detection of more subtle IBS activity contributing to task performance (Boas et al., 2001; Scholkmann et al., 2014). This discrepancy underscores HD-DOT’s advantage in capturing behaviorally meaningful neural signals, offering a more reliable framework for future hyperscanning research.

Despite the promising results, it is important to acknowledge several limitations of this study. As a preliminary study for proof-of-concept purpose, the focus on a specific type of collaborative task (e.g., tangram puzzles) may limit the generalizability of the findings to other forms of collaboration. Future studies could incorporate multimodal tasks (e.g., face-to-face dialogue or competitive games) to test the versatility of HD-DOT. Additionally, the relatively small sample size may have influenced the statistical power of the study. Increasing the sample size in future investigations would enhance the reliability and robustness of the findings. Finally, due to the limited optode number, the current HD-DOT montage used restricted optode density, which constrains our ability to map whole-brain IBS dynamics and likely underestimated the true extent of collaborative neural synchronization. Future implementations may prioritize expanded sensor density to capture distributed social brain networks.

## 5. Conclusion

By rigorously comparing HD-DOT and fNIRS, the present study advances hyperscanning methodology and deepens our understanding of the neural mechanisms driving social collaboration. The enhanced resolution of HD-DOT not only validates prior fNIRS findings but also uncovers novel IBS patterns, bridging a critical gap in social neuroscience. As optical neuroimaging evolves, HD-DOT promises to illuminate the neural underpinnings of human interaction with unprecedented clarity, offering new avenues for both basic research and clinical translation.

## CRediT authorship contribution statement

**Shuo Guan:** Conceptualization, Methodology, Formal analysis, Investigation, Writing-original draft, Visualization, Writing-review & editing. **Yuhang Li:** Methodology, Formal analysis, Writing-review & editing. **Yingbo Geng:** Methodology, Formal analysis, Writing-review & editing. **Dongyun Li:** Investigation, Resources, Writing-review & editing. **Qiong Xu:** Investigation, Resources, Writing-review & editing. **Dalin Yang:** Investigation, Resources, Software, Writing-review & editing. **Yingchun Zhang:** Validation, Formal analysis, Writing-review & editing. **Rihui Li:** Writing-review & editing, Resources, Methodology, Funding acquisition, Conceptualization, Project administration.

## Funding

This research was funded by the National Natural Science Foundation of China (82301743), the Science and Technology Development Fund of the Macao SAR (0010/2023/ITP1, 0016/2024/RIB1), Guangdong Provincial Natural Science Foundation (2025A1515010539), and the University of Macau (SRG2023–00015-ICI, MYRG-GRG2024-00296-ICI, and MYRG-CRG2024-00022-ICI).

## Declaration of competing interest

The authors declare that they have no known competing financial interests or personal relationships that could have appeared to influence the work reported in this paper.

## Ethics statement

All study protocols were conducted following the Declaration of Helsinki and were approved by the Institutional Review Boards at the University of Macau. Informed consent was obtained from each participant before the study.

## Data and code availability statement

The datasets used and analyzed during the current study are available from the corresponding author on reasonable request.

## Notes

### Competing Interest Statement

The authors have declared no competing interest.

